# MetaKSSD: Boosting the Scalability of Reference Taxonomic Marker Database and the Performance of Metagenomic Profiling Using Sketch Operations

**DOI:** 10.1101/2024.06.21.600011

**Authors:** Huiguang Yi, Xiaoxin Lu, Qing Chang

## Abstract

The rapid increase in genomes and metagenomic data presents major scalability and efficiency challenges for current metagenomic profilers. In response, we introduce MetaKSSD, which redefines reference taxonomic marker database (MarkerDB) construction and metagenomic profiling using sketch operations, offering efficiency improvements by orders of magnitude. MetaKSSD encompasses 85,202 species in its MarkerDB using just 0.17GB of storage and profiles 10GB of data within seconds, utilizing only 0.5GB of memory. Extensive benchmarking experiments demonstrated that MetaKSSD is among the top-performing profilers across various metrics. In a microbiome-phenotype association study, MetaKSSD identified significantly more effective associations than MetaPhlAn4. We profiled 382,016 metagenomic runs using MetaKSSD, conducted extensive sample clustering analyses, and suggested potential yet-to-be-discovered niches. Additionally, we developed functionality in MetaKSSD for instantaneous searching among large-scale profiles. The client-server architecture of MetaKSSD allows the swift transmission of metagenome sketches over the network and enables real-time online metagenomic analysis, facilitating use by non-expert users.

## 1. Introduction

Metagenomic profiling, which involves estimating the taxonomic composition and abundance of microbiome samples using whole genome sequencing (WGS) data, is a fundamental step in microbiome-related studies. These studies include microbiome-phenotype association studies^1–6^, microbiome-based disease prediction^7,8^, examining microbiome changes with age^9,10^, and microbiome co-occurrence and co-abundance studies^11–13^. Typically, a metagenomic profiling method (also called a profiler) estimates taxa abundances by mapping reads or *k*-mers from sample data to a pre-built reference taxonomic marker database (referred to as MarkerDB)^14,15^.

MarkerDB is normally constructed from genomes. Different profilers have different methods for processing genomes, making MarkerDB from one profiler incompatible with another. Most profilers, such as MetaPhlAn^9,16^ and mOTUs^17–19^, do not support custom MarkerDB construction. However, a few profilers, like Bracken^20^, do support it. The comprehensiveness of the MarkerDB is crucial, as a profiler can only detect taxon that exist in its MarkerDB. To maintain competitive performance, a profiler needs to update its MarkerDB periodically to cover the taxonomic markers from as diverse genomes as possible, especially metagenome-assembled genomes (MAGs)^21–23^. However, low-quality genomes, such as potentially misassembled ones, should be excluded. The taxonomic annotation for genomes is also crucial. There are two systems for taxonomic annotation: NCBI taxonomy^24^ and Genome Taxonomy Database (GTDB) taxonomy. NCBI taxonomy is the traditional taxonomy system adopted by most profilers^25,26^, whereas GTDB is newer system modified from NCBI taxonomy to provide rank-normalized, phylogenetically consistent taxonomy^27–30^.

In the past decade, more than 20 metagenomic profilers have emerged, each with its strengths at the time of release^14,25,26^. However, as the number of genomes grows exponentially^31^, most profilers face challenges in MarkerDB scalability. Bracken2^20^, a profiler that relies on the reads’ classifier Kraken2^32,33^, is perhaps an early victim of this trend, as its standard MarkerDB has grown to about 80GB^34^. Such a size poses difficulties for downloading in poor network conditions and running with limited memory resources. The predominant profiler, MetaPhlAn4^9^, has a MarkerDB of 20 GB, which has increased fivefold since its last iteration^16^ and will soon face similar issues. An exception is mOTUs^17–19^, whose MarkerDB is only 3.5GB and grows slowly with iterations.

With the continually dropping cost of sequencing, rapid and large-scale metagenomic profiling is also a trend. Bracken2, one of the fastest profilers revealed in the second round of the Critical Assessment of Metagenome Interpretation (CAMI2), requires about 0.66 hours to profile 100GB of data^25^. However, the data from a medium-scale microbiome sequencing project can easily surpass 5TB, requiring significant time for merely profiling. Furthermore, there are over 500,000 WGS metagenomic runs (total size estimated to be over 3PB) in the NCBI Short Read Archive (SRA)^35,36^ as of this study. Metagenomic profiling at this scale is necessary in the near future for constructing large models^37^ for microbiome data (Discussion). However, this task is far beyond the capacities of existing profilers.

Metagenomic profilers have a very wide user base; however, a significant proportion of potential users, such as clinical or wet lab scientists, may not be accustomed to command-line, which is the primary interface for most metagenomic profilers. Alternatively, they switch to online platforms such as Galaxy^38^ or One Codex^39^ for metagenomic analysis. However, in addition to potential data security problems, uploading sequencing data can be time-consuming and frequently fail, especially with limited bandwidth.

To meet these challenges, we present MetaKSSD, a metagenomic profiler developed upon the genome sketching technique K-mer Substring Space Decomposition (KSSD). KSSD is able to downsample a (meta)genome to a subset of *k*-mers (also called a sketch) with a size reduction of thousands of times, while maintaining the ability to accurately estimate metagenome containment using sketches^40^. KSSD uniquely enables lossless sketch operations such as union, intersection, and subtraction, which are fundamental for various sketch-based analyses^40^. Building upon these basic functionalities, MetaKSSD additionally introduces a feature that tracks *k*-mer counts within the sketch, and redefines the processes of MarkerDB construction and metagenomic profiling as a series of sketch operations (Results).

We demonstrate that MetaKSSD not only offers excellent MarkerDB scalability but also enables large-scale metagenomic profiling with ease. More importantly, the transmission of sample sketches over the Internet is swift, regardless of bandwidth limitations. In client-server mode, MetaKSSD realizes real-time online metagenomic profiling and instantaneous searches across hundreds of thousands of profiles for similar ones. This opens the door to a technique we call “sketch communications”, allowing users to search, compare, or share their sketches and profiles with others immediately (Discussion).

## 2. Results

### 2.1 Overview of MetaKSSD’s algorithm and main features

MetaKSSD redefines MarkerDB construction and metagenomic profiling using sketch operations, including union, intersection, and subtraction.

- **MarkerDB Construction**: Genomes are first sketched and organized by species (Fig. 1a). Inspired by the ‘pangenome’ concept^41^, a ‘pangenome sketch’ for each species is generated by uniting all sketches from that species. Species-specific markers are then created by subtracting the union of pangenome sketches of other species. Finally, all species-specific markers are consolidated into the MarkerDB (Fig. 1b, Methods).
- **Metagenomic Profiling**: For each sample, a sketch is created tracking *k*-mer counts. Summary statistics are then calculated from the counts of the *k*-mers overlapped with the species-specific markers in MarkerDB for each species, which could be normalized to determine the relative abundance (Fig. 1c, Methods).

**Fig. 1.**
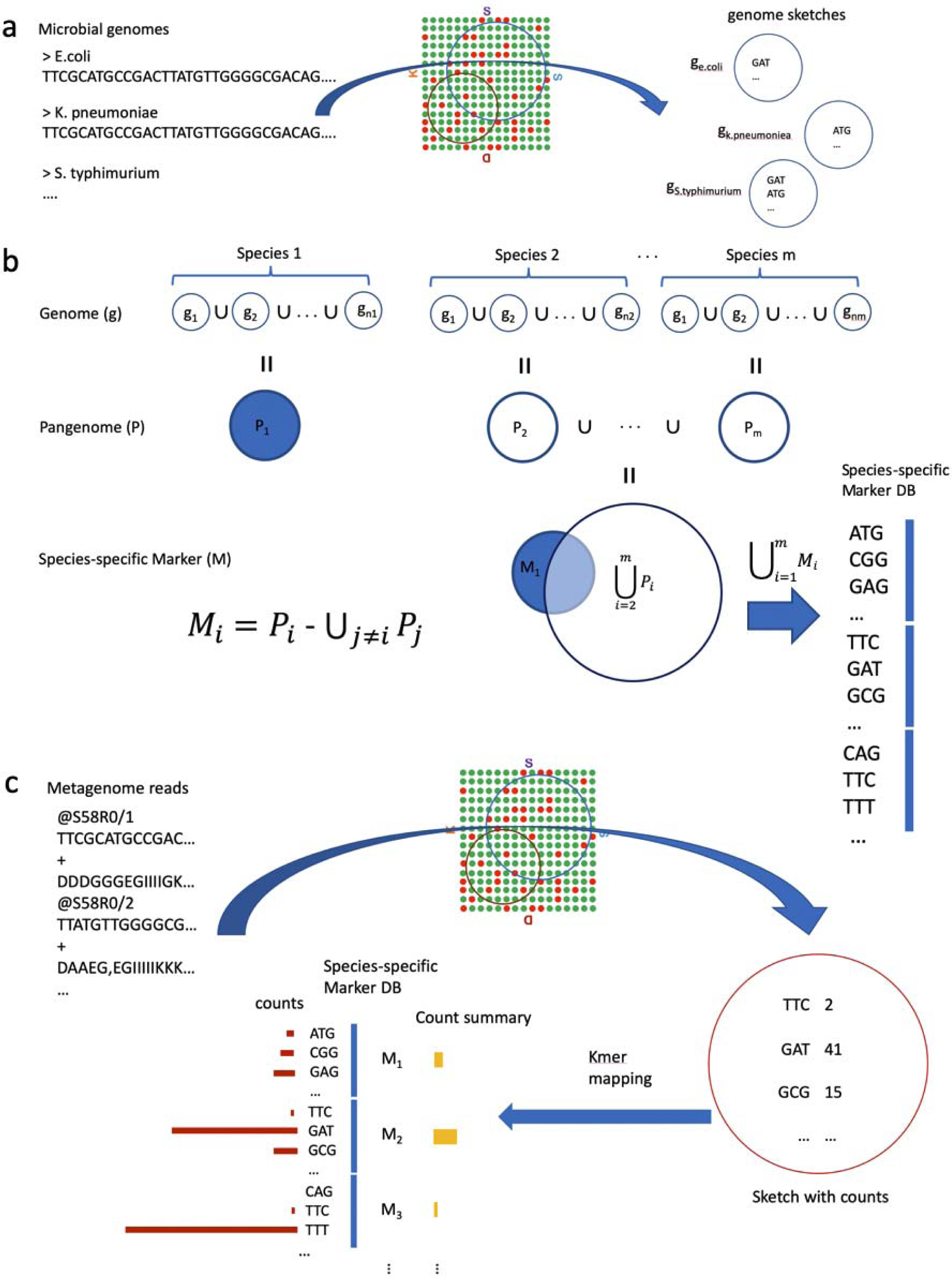
MetaKSSD algorithm overview. (a) Genomes are sketched and organized by species. (b) MarkerDB construction: For each species *i* (*i*=*1*..*m*, where *m* is the total number of species), the union sketch of all the sketches from species *i* is taken as its pangenome sketch *P_i_*. The species-specific marker *M_i_* is derived by subtracting from *P_i_* the union of all other pangenome sketches. All species-specific markers are indexed to build the MarkerDB. (c) Metagenomic profiling: The sample data is first sketched with *k*-mer counts tracked. The sketch is then overlapped with species-specific markers in the MarkerDB, and the abundance for each species is estimated using the summary statistics of those overlapped *k*-mer counts.

Constructed using sketches, MetaKSSD’s MarkerDB is orders of magnitude smaller than those constructed using whole genomes. It spans all 85,202 species of the Genome Taxonomy Database release 08-RS214 (GTDBr214), far surpassing MetaPhlAn4 and mOTUs3 (Fig. 2a). However, it requires only 0.17GB of storage for MarkerDB and 0.5GB of memory for profiling, vastly outperforming MetaPhlAn3 in both metrics (Fig. 2b). Therefore, MetaKSSD achieves a MarkerDB scalability, defined by the ratio of species number to storage usage, of 501,628 species per GB, which is 53 times higher than its closest competitor, mOTUs3 (9,429 species per GB).

**Fig. 2.**
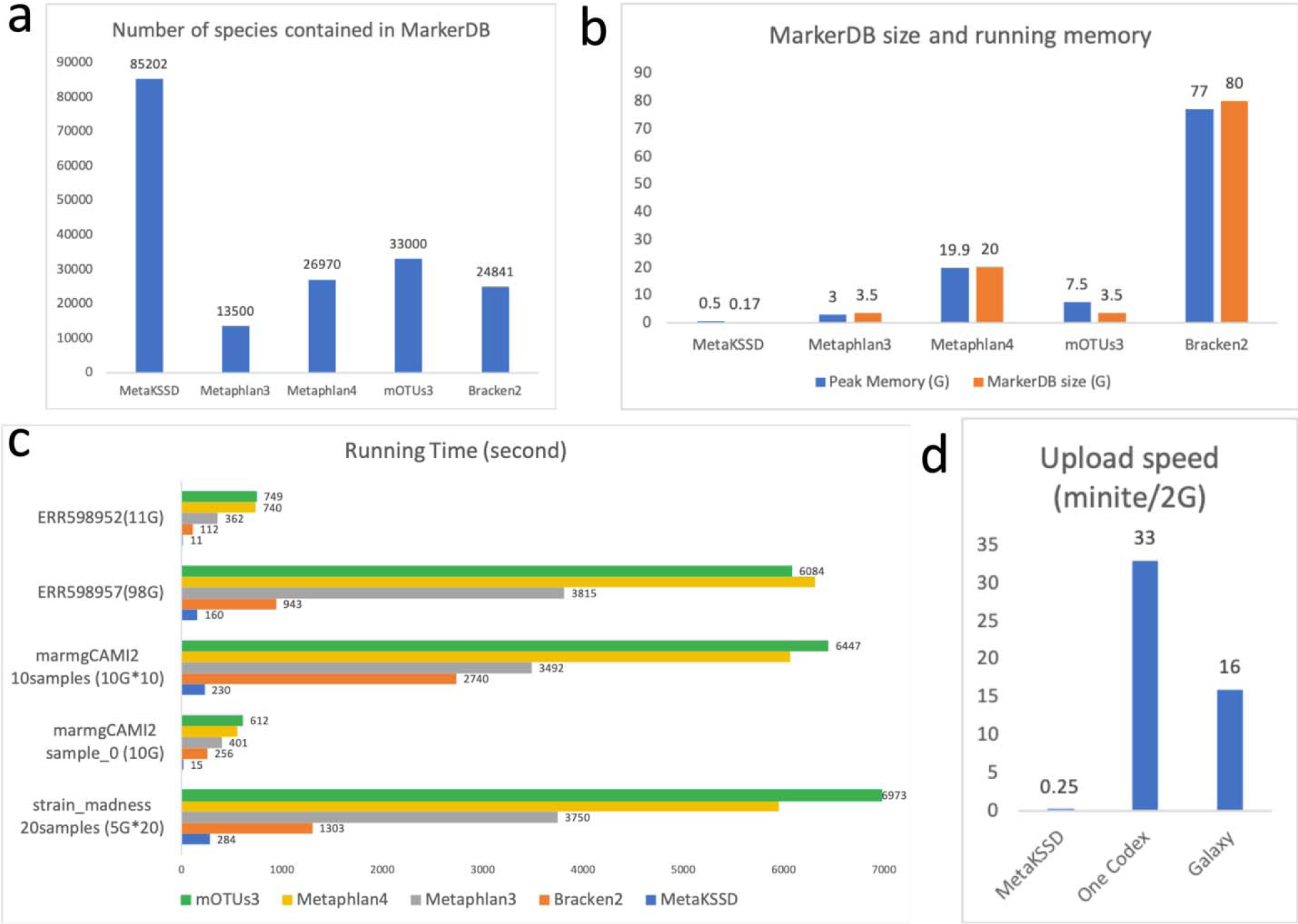
MetaKSSD quantifies more species while using fewer computational resources. Profiling speed testing (c) was conducted on an Amazon Web Services (AWS) c6a.24xlarge EC2 instance, featuring high I/O storage and utilizing 30 threads. Upload speed testing (d) was conducted on a MacBook Pro with a 40Mbps bandwidth

MetaKSSD operates on sketches, making the process of metagenomic profiling exceptionally fast. For instance, it profiled an 11GB metagenomic dataset (ERR598952) in just 11 seconds, which was 10 times faster than Bracken2, the fastest profiler from CAMI2. Additional tests demonstrated that MetaKSSD was 4.6 to 17 times faster than Bracken2, which was already significantly faster than other state-of-the-art profilers (Fig. 2c). As will be demonstrated later, MetaKSSD’s remarkable computational efficiency enables unprecedentedly large-scale metagenomic profiling.

MetaKSSD supports both command-line mode and a client-server architecture, enabling local sketching and server-based analysis (Code Availability). With a 4,096-fold dimensionality reduction, the typical metagenome sketch size is under 2 MB, allowing swift transmission over the network. Tests showed that the MetaKSSD client took 15 seconds to sketch a 2GB metagenome and 30 seconds for a full analysis, significantly outperforming Galaxy and One Codex (Fig. 2d).

In principle, the significant size reduction achieved by MetaKSSD does not compromise profiling quality. MetaKSSD estimates species abundance using read coverages on species-specific *k*-mers that are sparsely and randomly sampled from the whole genome. Each species in the 0.17GB MarkerDB occupies 1,820 bytes on average, equivalent to 455 species-specific *k*-mers. This is sufficient for abundance estimation except in rare cases of highly strain-diverse samples (Discussion).

### 2.2 Benchmarking MetaKSSD

Utilizing the Open-community Profiling Assessment tooL (OPAL)^42^, we conducted a comprehensive evaluation of MetaKSSD against the top performers identified in the CAMI2 taxonomic profiling challenges^25,26^. We focused on their latest iterations, including MetaPhlAn3^16^, MetaPhlAn4^9^, mOTUs3^17^, and Bracken2.8^20,32^ (by speed). These profilers were first benchmarked on four synthetic metagenome datasets from CAMI2^25^, representing diverse environmental contexts: Mouse gut (*n* = 49 samples), Rhizosphere (*n* = 21 samples), Marine (*n* = 10 samples), and Strain_madness (*n* = 100 samples) (Data Availability). Other profilers previously benchmarked on these datasets in CAMI2 were also included to provide a broader landscape for comparison (Data Availability). MetaKSSD was benchmarked across various parameter settings of *K* (the half-length of *k*-mer) and *S* (the minimum number of overlapped *k*-mers between a metagenome sketch and a MarkerDB species), while keeping *L* = 3 fixed (representing 4,096-fold dimensionality reduction). Since MetaKSSD used GTDB taxonomy, whereas other profilers and the ground truth used NCBI taxonomy, MetaKSSD’s results were converted to NCBI taxonomy to ensure comparability (Methods).

The evaluation considered four OPAL metrics at the species level:

1. **Purity (Specificity)**: The percentage of correctly predicted species among all predicted species in a metagenome sample.
2. **Completeness (Sensitivity)**: The percentage of correctly predicted species among all species in a metagenome sample.
3. **L1 Norm Error**: The estimation accuracy of species abundance.
4. **Weighted Unifrac Error**: The estimation accuracy of abundance across all taxonomic ranks.

The evaluation showed that no single profiler outperformed all others across all datasets, but the top performers consistently emerged from the MetaKSSD, MetaPhlAn, or mOTUs families. The sole exception was Dudes^43^, which achieved the highest purity in the ‘Strain_madness’ dataset. Notably, MetaKSSD excelled in the ‘Mouse_gut’ dataset, ranking highest in both purity and completeness (e.g., MetaKSSD.L3K11S48 with purity: 0.87±0.01, completeness: 0.976±0.003). MetaKSSD also ranked among the top in taxa abundance estimation accuracy (L1 norm error: 0.448±0.04, weighted Unifrac error: 1.24±0.0789), comparable to MetaPhlAn4 (L1 norm error: 0.412±0.0427, weighted Unifrac error: 1.19±0.0795). It also achieved the highest abundance accuracy in the ‘Rhizosphere’ dataset and was a top-tier profiler in the ‘Marine’ dataset. However, MetaKSSD showed only mediocre performance in the ‘Strain_madness’ dataset (Discussion) (Fig. 3).

**Fig. 3.**
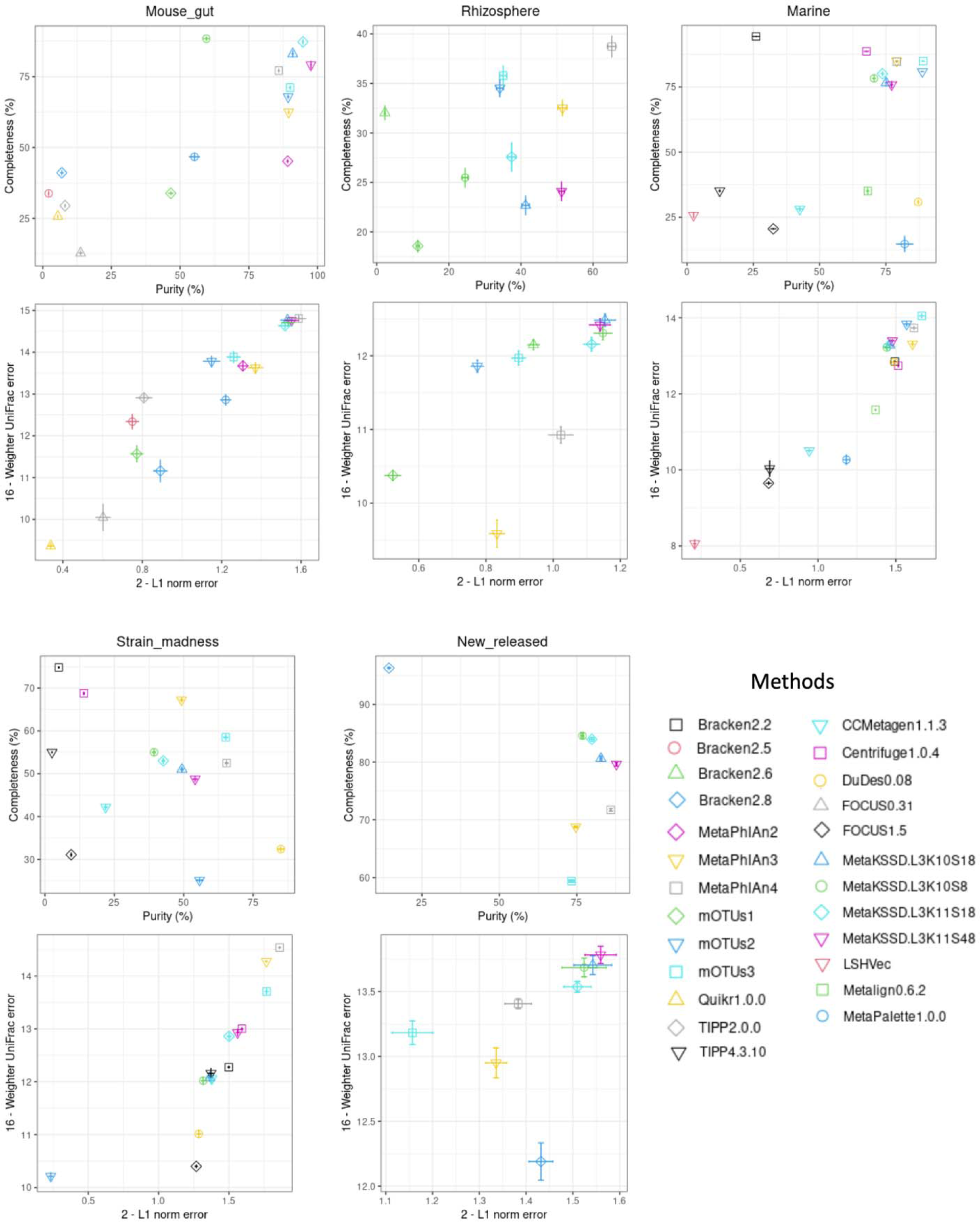
Performance comparison of MetaKSSD and other profilers. MetaKSSD, with different parameter settings, and various versions of other metagenomic profilers were benchmarked across five datasets: Mouse gut (*n* = 49 samples), Rhizosphere (*n* = 21 samples), Marine (*n* = 10 samples), Strain_madness (*n* = 100 samples), and the New_released dataset (*n* = 5 samples). Performance metrics include purity to completeness (the top row of each dataset) and l1 norm error to weighted unifrac error (the bottom row of each dataset). Purity measures the ratio of correctly predicted taxa to all predicted taxa, and completeness measures the ratio of correctly predicted taxa to all taxa present, both ranging from 0 (worst) to 1 (best). L1 norm error is the sum of absolute differences between true and predicted abundances, ranging from 0 (perfect) to 2 (totally incorrect). Weighted unifrac error measures the similarity between true and predicted abundances at all taxonomic ranks, ranging from 0 (high similarity) to 16 (low similarity). The width and length of the cross represents the standard deviations of the corresponding metric for each data point.

MetaKSSD’s performance on the ‘Rhizosphere’, ‘Marine’, and ‘Strain_madness’ datasets may have been underestimated because their source genomes were anonymous, preventing precise GTDB to NCBI taxonomy conversion (Methods). Therefore, errors could arise from the differences in taxonomy systems, rather than genuine inaccuracies. In contrast, the ‘Mouse_gut’ dataset provided source genome IDs, enabling precise taxonomy conversion and improved performance.

Later-developed profilers generally outperformed older ones, as seen with MetaPhlAn4 and mOTUs3 versus MetaPhlAn2,3 and mOTUs1,2. These improvements might be due to the inclusion of CAMI2 source genomes in later-built MarkerDBs rather than methodological innovations, as noted in previous benchmarking studies^14,15^. To address this, we constructed five synthetic metagenome samples (referred to as the ‘New released’ dataset) using genomes absent from all MarkerDBs (Tab. S1) for further benchmarking (Methods). On the ‘New released’ dataset, MetaKSSD outperformed other top profilers in almost all metrics (Fig. 3), with the sole exception being Bracken2.8, which achieved the highest completeness but the lowest purity. This may be because Bracken2.8 used all taxonomically informative *k*-mers from whole genomes^33^, while others used only quality-controlled marker genes (MetaPhlAn and mOTUs families) or sketches of markers (MetaKSSD).

In addition to species-level evaluations, MetaKSSD achieved similar performance across other taxonomic ranks (Figs. S1-S5). Overall, although no single metagenomic profiler was the best for all situations, MetaKSSD emerged as one of the top performers in most cases.

### 2.3 MetaKSSD improves microbiome-phenotype association study

The unprecedentedly comprehensive MarkerDB of MetaKSSD presumably allows for more refined metagenomic profiling. However, this advantage has not fully materialized due to the GTDB to NCBI taxonomy conversion in the benchmarking experiments described above. To explore its capabilities with the native GTDB taxonomy system, MetaKSSD was applied to a microbiome-phenotype association study within the BGInature2012 cohort^44^ and compared with MetaPhlAn4, the predominant metagenomic profiler for microbiome-phenotype association studies^1,45,46^.

The BGInature2012 cohort was selected for its well-characterized and fully open-access nature (Data Availability), unlike other cohorts that typically require an application and review process. Its datasets include stool microbiome WGS data and related metadata from 368 Chinese individuals. The metadata encompass gender, age, and 12 physiological phenotypes: diabetic status, body mass index (BMI), fasting blood glucose (FBG), systolic blood pressure (SBP), diastolic blood pressure (DBP), fasting serum insulin (FINS), fasting serum C-peptide (FCP), glycosylated hemoglobin HbA1c (HbA1c), triglycerides (TG), total cholesterol (TCHO), high-density lipoprotein (HDL), and low-density lipoprotein (LDL). Based on the species abundance profiles generated by both MetaKSSD and MetaPhlAn4 for all 368 samples, the species significantly associated with each of these phenotypes were identified (Methods).

Out of the 12 phenotypes examined, MetaKSSD identified associations for 9 phenotypes, while MetaPhlAn4 identified associations for 7 phenotypes, missing findings in FBG and LDL. Neither method found associations in the three phenotypes—DBP, TCHO, and BMI—thus, they were excluded from our comparisons. Interestingly, associations emerged upon converting BMI to obesity status based on Chinese obesity criteria (BMI ≥ 28). Therefore, BMI was replaced by obesity status, resulting in a total of 10 phenotypes in the comparisons. MetaKSSD consistently identified more associations than MetaPhlAn4 across the 10 phenotypes (Fig. 4a, b). In total, MetaKSSD identified 49 associated species, comprising 23 knowns and 26 unknowns (yet-to-be cultivated or named, detailed in Methods), whereas MetaPhlAn4 identified 28, consisting of 23 knowns and 5 unknowns (Tab. S2). Therefore, the increased number of associated species identified by MetaKSSD is mainly contributed by unknown species, a trend that persists across all phenotypes (Fig. 4b).

**Fig. 4.**
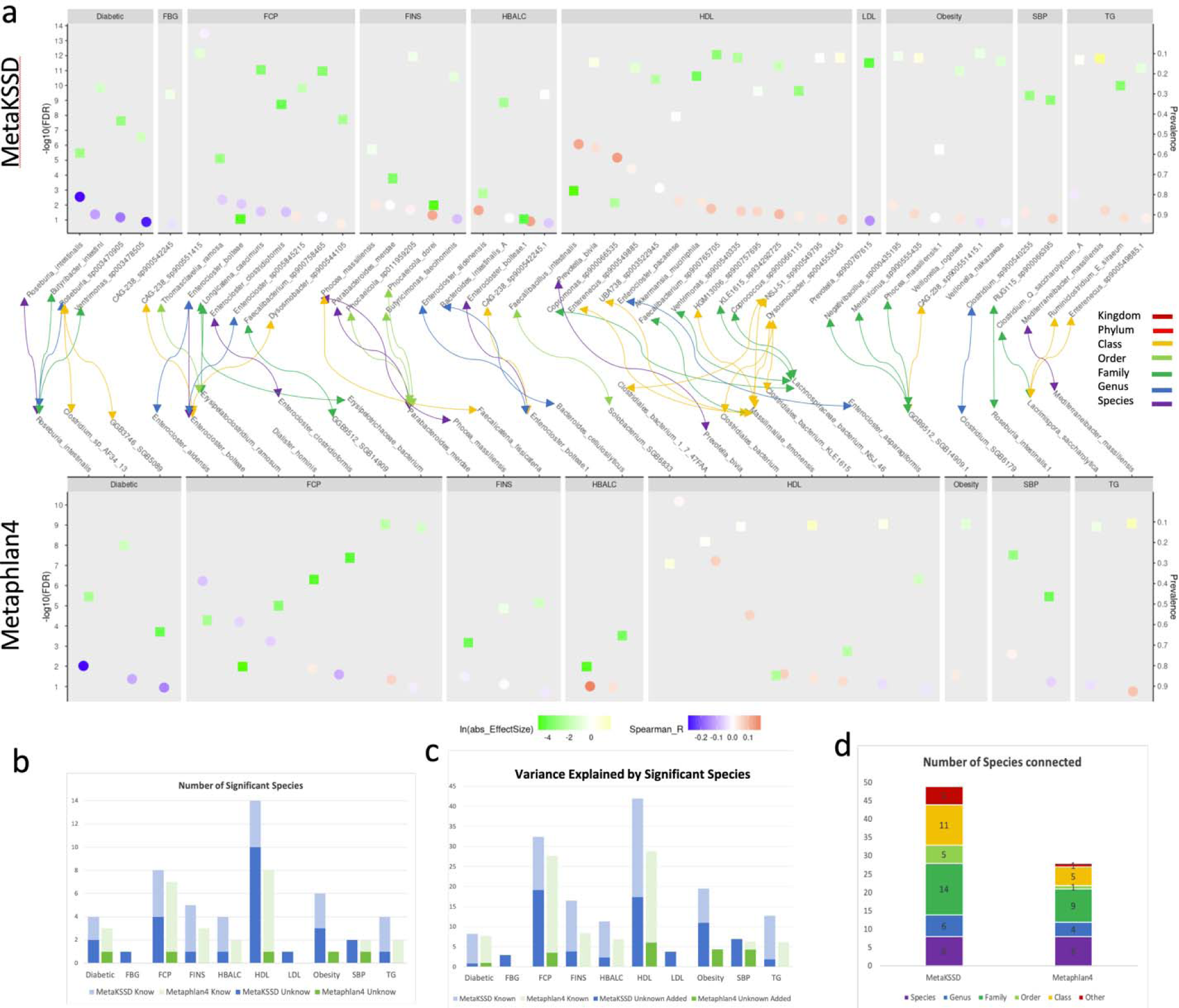
Comparison of MetaKSSD and MetaPhlAn4 for microbiome-phenotype association study. All 368 Chinese stool samples from the BGInature2012 cohort^44^ were profiled using both MetaKSSD and MetaPhlAn4, identifying bacterial species associated with 10 phenotypes: diabetic status, fasting blood glucose (FBG), fasting serum C-peptide (FCP), fasting serum insulin (FINS), glycosylated hemoglobin HbAlc (HBALC), high-density lipoprotein (HDL), low-density lipoprotein (LDL), obesity status, systolic blood pressure (SBP), and triglyceride (TG). (a) Negative log-transformed false discovery rate (FDR, circle dots, left y-axis) and prevalence (square dots, right y-axis) of the bacterial species (x-labels) identified by MetaKSSD and MetaPhlAn4 for each phenotype. A line connects a MetaKSSD species with its closest MetaPhlAn4 species if they belong to the same class, with the color of the line indicating their lowest common ancestry rank. Species name prefixed with the pattern “sp[0-9]{9}” and “SGB” are unknown species for MetaKSSD and MetaPhlAn4, respectively; otherwise, they are known species. Comparison of MetaKSSD and MetaPhlAn4 with regarding (b) the number of significant species identified and (c) the percentage of variance explained. (d) Number of species connected in (a) by different taxonomic ranks.

The analysis of phenotypic variances showed that MetaKSSD species consistently explained a higher percentage of variance than MetaPhlAn4 species across all phenotypes (Fig. 4c, Methods), affirming the effectiveness of these increased species of MetaKSSD. By subtracting the percentage of variance explained by known species from the total percentage of variance explained, the “unknown added” was calculated, representing the percentage of variance solely explained by unknown species, excluding the part explained by both known and unknown species. The “unknown added” constituted a substantial proportion of the total variance explained when unknown species were present (Fig. 4b, c), affirming their effectiveness. Compared to MetaPhlAn4, MetaKSSD’s unknown species explained a significantly higher percentage of variance across most phenotypes, except for diabetic status, where there was a tie (Fig. 4c).

There are 8 common species detected by both profilers, accounting for only 16% (8 out of 49 species) and 29% (8 out of 28) of the species identified by MetaKSSD and MetaPhlAn4, respectively. However, extensive connections between the two profilers’ results were observed at higher taxonomic ranks (Fig. 4a, Tab. S3). Specifically, ninety percent of MetaKSSD species (44 out of 49) have a MetaPhlAn4 counterpart from the same class, while vice versa, 96% (27 out of 28) also hold true (Fig. 4d). This observation aligns with the fact that the GTDB taxonomy (used by MetaKSSD) is much more consistent with the NCBI taxonomy (used by MetaPhlAn4) at the class rank than at the species rank^27–30^. This consistency at higher taxonomic ranks indicates a strong agreement between the results of the two profilers and supports the overall reliability of both profilers’ results despite differences at the species level.

More than 40% (20 out of 49) of the findings presented in this study are supported by prior research (see Tab. S3 for references). For instance, both MetaKSSD and MetaPhlAn4 identified *Roseburia intestinalis* as negatively correlated with diabetes, aligning with findings from previous studies^47,48^. Similarly, MetaKSSD detected an association between *Parabacteroides merdae* and fasting serum insulin, which was also reported by Qiao et al.^49^. These extensive agreements underscore the reliability of our microbiome-phenotype association identification approach.

In summary, the results highlight that MetaKSSD enhances microbiome-phenotype association studies by identifying a significantly larger number of species associated with phenotypes. Importantly, many of these associations involve unknown species that were not previously recognized by earlier studies, underscoring the potential of MetaKSSD to uncover novel insights in microbiome research.

### 2.4 Large-scale metagenomic profiling and profile searching with MetaKSSD

MetaKSSD was applied to analyze metagenomic data publicly available in NCBI SRA. Out of the 517,829 WGS metagenomic runs retrieved, 382,016 passed our quality control and were successfully sketched and profiled, spanning 312,664 samples,16,398 projects and 2,025 environments (detailed in Methods; see Tab. S4 for the runs information and Data Availability for all sketches and profiles). Summarizing all the profiling results revealed that the number of samples versus the number of detected species (and genera) follows a long-tail distribution (Fig. S6). Specifically, thresholding the number of MarkerDB overlapping *k*-mer at 18, 97% of samples contain fewer than 1,000 species. However, 0.14% of samples contain over 10,000 species, which are mostly from soil, rhizosphere or wetland (Tab. S4). The remarkably high species diversities observed in these environments are consistent with findings from previous studies^50,51^.

Fifty-three commonly studied metagenomic environments were identified (detailed in Methods; see Tab. S5 for the list), accounting for 49% of the total samples. In these environments, the ‘bovine gut’ exhibited the highest median species number per sample (median *m* = 671), significantly exceeding those of the ‘mouse gut’ (*m* = 164) and ‘human gut’ (*m* = 145). Considering that a larger proportion of ‘bovine gut’ *k*-mers are absent from GTDBr214, the extent of this excess is likely underestimated. While ‘wetland’ included the sample with the largest species number (24,554 species for sample SAMN20864071), it had a relatively lower median species number (*m* = 327.5) (*S* = 18, Fig. 5a). Similar environment rankings were also observed when using a threshold of *S* = 48 or at the genus level (Fig. S7).

**Fig. 5.**
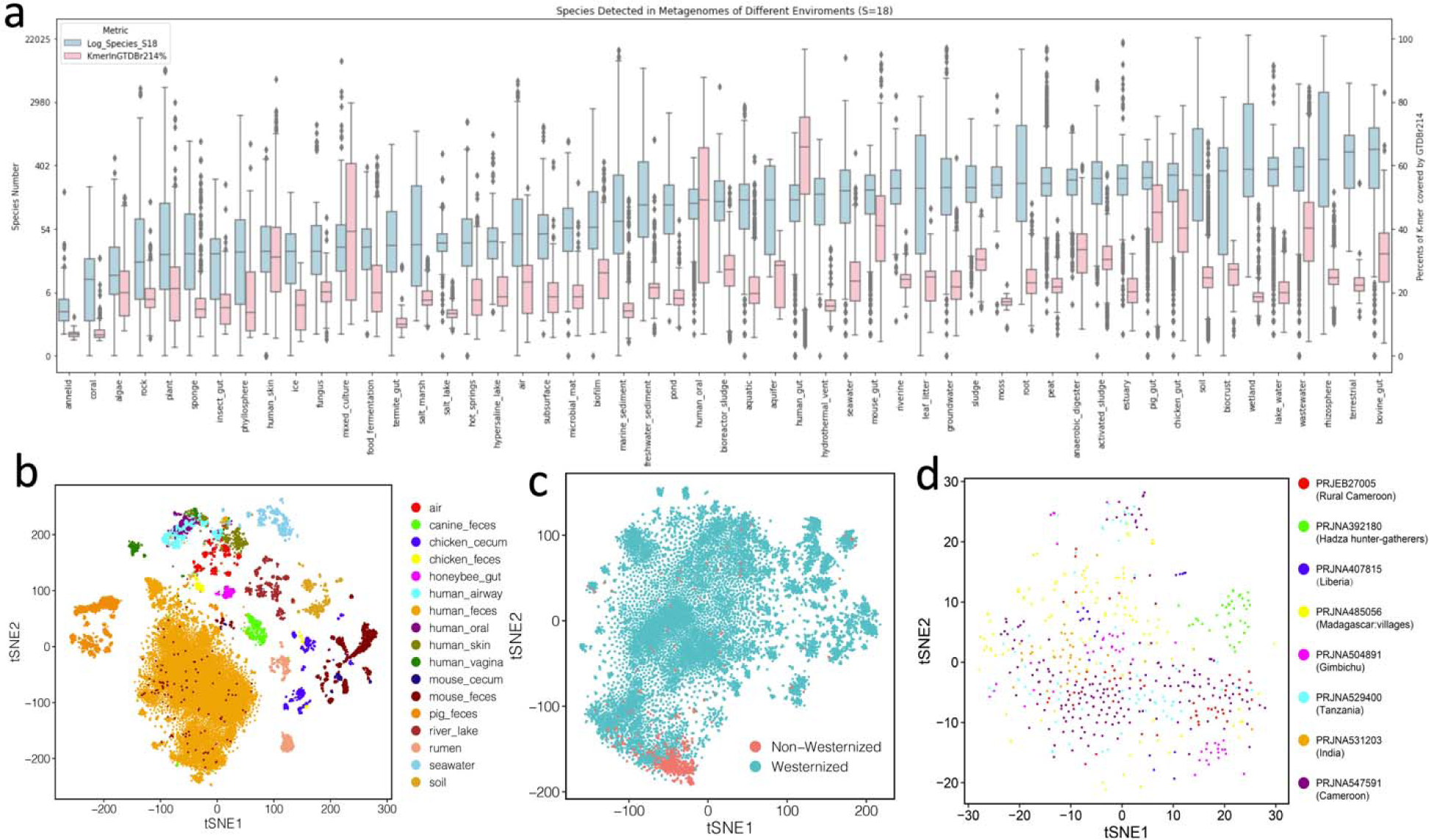
Large-scale metagenomic profiling. (a) Species number (left y-axis) detected in different environments alongside *k*-mer percentages covered by GTDBr214 genomes (right y-axis). The species number for each run from the 53 commonly studied metagenomic environments (Tab. S5) was determined from its MetaKSSD profile, using a MarkerDB *k*-mer overlap threshold of *S* = 18. The percentage of *k*-mers (*k* = 22) covered by GTDBr214 was estimated by the proportion of *k*-mers in a run sketch that overlapped with the union sketch of all GTDBr214 genomes. The species number and *k*-mer percentage of a sample are the averages of the species numbers and *k*-mer percentages of all its runs. Environments were ranked by the sample median of species number. (b)-(d) T-SNE analysis of 151 well-characterized projects (Tab. S7), lifestyle-related projects (Tab. S8), and non-westernized samples, respectively.

All detected species, after controlling for suspicious findings (Methods), were ranked by the number of environments in which they are present. The top species, as expected, are well-characterized species such as *Escherichia coli*, *Klebsiella pneumoniae*, and *Salmonella enterica* (Fig. S8). In total 17,742 environment-specific species were identified, spanning all the fifty-three environments (Tab. S6), which may serve as an important reference for prokaryote habitats studies^52^. These results provide a general characterization of the landscape of existing metagenomic data, which has been difficult to achieve using conventional metagenomic profilers.

The application of t-SNE (t-distributed Stochastic Neighbor Embedding) analysis on a subset of 151 well-characterized projects, encompassing 30,592 samples across17 different environments (see Tab. S7 for accessions and Methods for enrollment criteria), revealed distinct clusters of samples based on their respective environments. Within specific environments, such as mouse feces and air (brown and red dots in Fig. 5b, respectively), sub-clusters were frequently observed. Further investigation indicated that these sub-clusters corresponded to specific projects (Fig. S9), suggesting potential influences from batch effects (i.e., technical cause) or the presence of project-specific, uncharacterized sub-environments (i.e., biological cause). To delve deeper into the primary cause of these sub-clusters, t-SNE analysis was conducted on a subset of human stool samples related to different lifestyles, which were well-characterized for their originating environments (Tab. S8, initially adopted by the MetaPhlAn4 work^9^). This analysis revealed a distinct separation between samples from westernized (16,669 samples, 60 projects) and non-westernized (560 samples, 8 projects) lifestyles, with the latter forming a clearly delineated cluster (Fig. 5c). These findings demonstrated similar discriminatory power in distinguishing between westernized and non-westernized clusters as observed in previous MetaPhlAn4 results^9^. Further exploration of non-westernized samples showed that samples from seven different projects were intermingled, indicating no significant project-specific batch effects. The only exception was PRJNA392180 (green dots in Fig. 5d), comprising 40 Hadza hunter-gatherer gut microbiome samples, which formed a distinct cluster. Previous studies using different datasets have supported the notion that Hadza hunter-gatherers exhibit a unique gut microbiome profile^53^. Taken together, these observations suggested that the sub-clusters identified using MetaKSSD were likely attributable to biologically meaningful, often uncharacterized sub-environments rather than batch effects.

To optimize computing efficiency, the abundance profiles were encoded as sparse vectors and saved as binary files. As a result, the entire abundance vector database, encompassing all 382,016 runs, required only 400MB of storage (Data Availability). Given an abundance vector, MetaKSSD can immediately identify the most similar vectors from the database using two optional distance (or similarity) measures between the query and each abundance vector in the database (Methods):

1. Normalized L1 norm (Eq. 1)
2. Cosine similarity (Eq. 2)

Both measures can effectively group samples by their originating environments using the multidimensional scaling (MDS) method, as demonstrated with the test abundance vectors (Fig. S10, Tab. S9). This supported the use of these two measures for profile searching. The abundance vector searching functionality is particularly useful in a client-server mode (Code Availability), enabling users without bioinformatics skills to easily explore similar profiles in the database for their samples (Discussion).

## 3. Discussion

The performance of a metagenomic profiler is limited by the comprehensiveness of its MarkerDB^14,15,54^. However, preparing and updating MarkerDB is often cumbersome and significantly lags behind the rapid release of new genomes^14,31,55^. This study proposes MetaKSSD as a swift and highly scalable approach for MarkerDB preparation, enabling the construction of a more comprehensive MarkerDB while using significantly fewer computational resources. With this more comprehensive MarkerDB, MetaKSSD enhanced microbiome-phenotype association studies compared to the current state-of-the-art methods.

Compared to existing profilers, MetaKSSD improved time and memory efficiency by orders of magnitude without compromising accuracy. These advancements facilitated large-scale metagenomic profiling and data-driven discoveries. For instance, the clustering of large-scale metagenome profiles revealed uncharacterized groups, suggesting potential yet-to-be-discovered niches, such as unknown lifestyles or health statuses. However, further sample surveys and experimental validations are essential to confirm these findings. Additionally, recent research has shown that a large ‘language’ model (LLM) trained on large-scale single-cell profiles can broadly benefit the field of single-cell biology^56^. With the exponential growth of metagenomic data^55^, a similar metagenomic LLM could be trained based on large-scale MetaKSSD profiles in the near future. In this analogy, a profile is a ‘sentence’ and species abundances are the ‘words’. Such a metagenomic LLM might be used for species abundance enhancement, imputation, batch effect correction, sample annotation, and other applications, as demonstrated in the example of single-cell LLM^56^.

Before profiling, MetaKSSD reduces metagenomic data to a sketch typically smaller than 2 MB, allowing it to be swiftly transmitted over the Internet regardless of the bandwidth limitations. This feature makes MetaKSSD (in client-server mode) the first real-time online metagenomic analysis platform. This platform is designed to assist a broad range of users in metagenomic analysis, including those with no bioinformatics skills, such as physicians or laboratory medicine specialists for clinical use^57^. Metagenomic analysis can be performed promptly by simply selecting and submitting FASTQ files in the processing window (see Code Availability for MetaKSSD clients). There is no technical barrier to extending this platform to support various types of omics data, such as genomic, transcriptomic, or epigenomic data, which would transform it into an omics sketches hub. Utilizing the established distance estimation functionalities for sketches^40^ and profiles (Methods), this platform would allow users to search, compare and share sketches and profiles promptly with each other. However, addressing the related privacy and data security concerns remains part of our future plans.

MetaKSSD still faces some limitations. It estimates species abundance using the average of the 98^th^ and 99^th^ percentiles of MarkerDB overlapped *k*-mer counts (Methods). This approach aims to provide a simplified estimation of species abundance when a sample potentially contains two or more strains of a species, without involving complex statistical modeling. Although this measurement performed well on most datasets (typically containing fewer than three strains per species), it significantly underperformed on the highly strain-diverse dataset ‘Strain-madness’ (containing more than 20 strains per species)^25^. Fortunately, species in real-world microbiomes are typically dominated by a single strain^58–61^, which justifies the application of MetaKSSD. MetaKSSD has also been tested on 16S rRNA data (using 16-fold dimensionality-reduction, L1K9) and long-reads data but showcased inferior performance compared to using WGS short reads data (not shown). Further improvements to MetaKSSD on these metagenomic data types are feasible but not deemed worthwhile, since metagenomic profiling using 16S rRNA data is already computationally efficient and long-reads metagenomic data is primarily used for assembly rather than profiling^62^.

## 4. Methods

### 4.1 MarkerDB construction

All 402,709 genomes from GTDBr214 were downloaded using the accessions provided in the metadata files “bac120_metadata_r214.tar.gz” and “ar53_metadata_r214.tar.gz”, accessible at https://data.gtdb.ecogenomic.org/releases/release214/214.0/. These genomes were sketched by MetaKSSD (Code Availability) using the following command:

‘‘‘

metakssd dist -L <L3K11.shuf> -o <L3K11_sketch> <all_genomes_Dir>

‘‘‘

where “L3K11.shuf” is the parameter file defining the dimensionality reduction level (L) as 3 and the *k*-mer length as 22. The dimensionality reduction rate is 16*^L^*, resulting in 4,096 when *L* = 3, and the *k*-mer length is twice the value of *K*, equating to 22 when *K* = 11. The resulting sketch of all the genomes was stored as “L3K11_sketch”.

The genome names within the sketch were printed line by line and directed to a file (e.g., “genome_name.txt”) using the following command:

‘‘‘

metakssd set -P <L3K11_sketch> > <genome_name.txt>

‘‘‘

Then, a grouping file (e.g., “group_name.txt”) was prepared as follows: The file “group_name.txt” should contain the same number of lines as “genome_name.txt”. Each line specifies the species name of the genome corresponding to the line in “genome_name.txt”. The format for each line should be: “ID species_name”, where the ID is a non-negative integer uniquely labeling the species_name. The species name of the genome can be found in the metadata files “<u>.*_metadata_r214.tar.gz</u>”. For example, “1 Escherichia_coli” represents the species name “Escherichia_coli” labeled with ID 1. The ID 0 is reserved for excluding genomes from the resulting sketch. Once the file “group_name.txt” is prepared, genomes can be grouped by their originating species using the following command:

‘‘‘

metakssd set -g <group_name.txt> -o <L3K11_PAN-sketch> <L3K11_sketch>

‘‘‘

Here, “L3K11_pan-sketch” represents the consolidated ‘pangenome’ sketches for all species (Fig. 1b).

Subsequently, the union sketch of all species-specific markers (e.g., “L3K11_union_sp-sketch”), was obtained using this command:

‘‘‘

metakssd set -q -o <L3K11_union_sp-sketch> <L3K11_pan-sketch>

‘‘‘

Finally, the MarkerDB (e.g., “markerdb_L3K11”) was generated by overlapping “L3K11_union_sp-sketch” with “L3K11_pan-sketch” using the following command:

‘‘‘

metakssd set -i <L3K11_union_sp-sketch> -o <markerdb_L3K11> <L3K11_pan-sketch>

‘‘‘

### 4.2 Metagenomic profiling and summary statistics of the *k*-mer counts

For metagenomic profiling, the microbiome sample (e.g., “sample1.fastq”) was first sketched with *k*-mer counts tracked using the following command:

‘‘‘

metakssd dist -L <L3K11.shuf> -A -o <sample1_sketch> <sample1.fastq>

‘‘‘

Here, the parameter file “L3K11.shuf” imposed a 4,096-fold dimensionality reduction for sketching.

Subsequently, summary statistics of *k*-mer counts (namely, read coverages on *k*-mers) for all identified species were generated with this command:

‘‘‘

metakssd composite -r <markerdb_L3K11> -q <sample1_sketch> > <species_coverage.tsv>

‘‘‘

“markerdb_L3K11” referred to the pre-computed MarkerDB (Methods section 4.1).

The resulting table, “species_coverage.tsv”, comprised seven columns: sample name, species name, number of overlapping *k*-mers, mean read coverage, average of the 98^th^ and 99^th^ percentiles of *k*-mer counts, median *k*-mer counts, and maximum *k*-mer counts. The fifth column (the average of the 98^th^ and 99^th^ percentiles of *k*-mer counts) was normalized to the relative abundances. This choice was made based on the performance of different summary statistics (Fig. S11), as well as the following considerations:

1. The microbiome may contain multiple strains within a single species, resulting in stacked read coverages for common *k*-mers. Therefore, the *k*-mer counts at higher percentiles provided a more accurate estimation of species abundance.
2. The top 1% of *k*-mer counts were likely attributed to repetitive regions and were therefore excluded to avoid potential biases.

Normalization was conducted using this command:

‘‘‘

perl possion.kssd2out.pl <species_coverage.tsv> <least_overlapped_*k*-mers > >

<species_relative_abundance_profile>

‘‘‘

Here, the “least_overlapped_*k*-mers” was an integer, defaulting to 18, to control false discoveries.

### 4.3 Simulation of the ‘New_released’ dataset and taxonomic systems conversion

The ‘New_released’ dataset was simulated using CAMISIM^63^, with the same settings as the <u>CAMI2 challenge datasets</u>, but used source genomes that were not present in any profiler’s MarkerDB. These source genomes were selected as follows: Complete genomes released after the latest MarkerDB (March 2022 for mOTUs3) were chosen from the latest <u>summary file</u> for bacteria genome assemblies. The genomes present in GTDBr214 (used by MetaKSSD) were explicitly excluded, ensuring the genomes enrolled were not present in any profiler’s MarkerDB. This step yielded 5,193 genomes. To prevent over-representation of common species such as *Salmonella enterica* and *Escherichia coli*, only species harboring 2 to 15 genomes out of the 5,193 were selected. Furthermore, only genomes from species with consistent taxonomy between NCBI and GTDBr214 were chosen. This selection process resulted in a total of 162 species, with one genome randomly selected from each (see Tab. S1 for accessions).

MetaKSSD used GTDBr214 taxonomy, while all other profilers and the ground truth used NCBI taxonomy. Therefore, the GTDBr214 to NCBI taxonomy conversion was performed on MetaKSSD results before comparison. The ‘New_released’ dataset was simulated using taxonomy-consistent genomes, so its taxonomy conversion involved a straightforward one-to-one mapping from GTDBr214 to NCBI species of the source genomes (Tab. S1). For the ‘Rhizosphere’, ‘Marine’, and ‘Strain_madness’ datasets, the source genomes were anonymous, and their GTDBr214 taxonomy could not be tracked. For these datasets, the GTDBr214 to NCBI species mapping was created based on the <u>taxonomies of all the GTDBr214 genomes</u> using the maximal supports rule: If a GTDBr214 species consists of genomes from multiple NCBI species, then the NCBI species with maximal number of genomes from this GTDBr214 species was mapped (see Tab. S10 for the mapping scheme created based on this rule). This conversion scheme can introduce errors if a source genome belongs to a minor NCBI species, potentially underestimating MetaKSSD’s performance on these datasets. For the ‘Mouse_gut’ dataset, which has all its source genomes’ accession IDs disclosed, a GTDBr214 to NCBI species mapping scheme was created and applied for taxonomy conversion for each sample based on the taxonomies of the sample’s source genomes.

### 4.4 Microbiome-phenotype association study

All the sequencing datasets of the BGInature2012 cohort^44^ were first downloaded (Data Availability). For metagenomic profiling, MetaPhlAn4 used default parameters and the MarkerDB vJan21 (the version used in its original paper^9^). MetaKSSD sketched these data using a 22-mer at a 4,096-fold dimensionality-reduction rate (with the parameter file “L3K11.shuf”). Since a spurious control measure was applied later, the MarkerDB overlapping threshold was not set here to retain as many species as possible.

After profiling, species relative abundances from individual profiles were compiled into a matrix, with rows representing species and columns representing sample IDs. To control for spurious species, those with a prevalence of less than 0.1 were removed, retaining only species with a non-zero abundance in more than 36 individuals.

All phenotypic measurements of the BGInature2012 cohort were obtained from the supplementary tables of its original paper^44^. Missing values in these measurements were imputed with the mean of the available data. One of these phenotypes, body mass index (BMI), was converted to obesity status (0 or 1) using Chinese obesity criteria (BMI ≥ 28)^64,65^.

Microbiome-phenotype associations were detected across all species-phenotype pairs using a linear model^66^ in R ^67^, including age and gender as covariates:

‘‘‘

Phenotype ∼ Age + Gender + Species Relative Abundance

‘‘‘

P-values of the associations were subjected to Benjamini-Hochberg correction^68^ to control for multiple testing. According to the conventions of MetaPhlAn4^6,9^, an association was considered significant at a false discovery rate (FDR) < 0.2.

Spearman correlation between phenotype and species was calculated using the ‘cor’ function^69^ in R. To calculate the percentage of phenotypic variance explained by the significantly associated species, a linear model^66^ was applied:

‘‘‘

fit <-lm (phenotype ∼ sp_1_ + sp_2_ + … + sp_n_)

‘‘‘

where the variable ‘phenotype’ was the phenotypic measurements, and sp_1_, sp_2_, …, sp_n_ were the relative abundances of the significantly associated species. The value ‘summary(fit)$r.squared’ was reported as the percentage of phenotypic variance explained. The percentage of variance explained by known species was calculated in the same way using only known species in the model. For MetaPhlAn4 results, known species were defined as species not prefixed with “_SGB”; for MetaKSSD results, known species were defined as species not prefixed with the pattern “_sp[0-9]{9}”; otherwise, they were considered known species, following the naming conventions in MetaPhlAn4^9^ and GTDB^27,29,30^.

### 4.5 Sketching of large-scale metagenomic data

All WGS metagenomic runs available as of Dec. 27, 2023, were obtained using the following search query executed at <u>NCBI SRA</u>:

‘‘‘

((((((Public[Access]) AND (“0000”[Modification Date] : “2023/12/27”[Modification Date])) AND “wgs”[Strategy]) AND “metagenomic”[Source] AND “cluster_public”[prop] AND “filetype fastq”[Properties])) AND “filetype fastq”[Filter]) AND “cloud s3”[Filter]

‘‘‘

The returned run information, totaling 517,829 runs, was saved in a file (e.g., “SraRunInfo.csv”) for downstream analysis.

Subsequently, undesired runs using sequencing technologies such as 16S rRNA, amplicon, nanopore, pacbio, and solid were excluded with the following command:

‘‘‘

awk -F ‘,’ ‘$5>2e8 && $5 > $8*1e6 && $13 == “WGS” && $14 ∼ /PCR|RANDOM|unspecified/ && $15 == “METAGENOMIC” && $19 !∼ /NANOPORE|PACBIO|SOLID/ ‘ SraRunInfo.csv |grep -iv ‘dbgap\|16S’ | csvcut -c1 > accession.list

‘‘‘

This filtering resulted in 420,708 runs. Runs were sketched in parallel on Amazon Web Services (AWS) instances for efficiency:

‘‘‘

nohup cat accession.list | parallel -j 16 “ aws s3 cp s3://sra-pub-run-odp/sra/{}/{} $TMPDIR/ && kssd221 dist -L <L3K11.shuf> -A -o <outdir> --pipecmd ‘fastq-dump --skip-technical -- split-spot -Z’ $TMPDIR/{} && rm $TMPDIR/{} “ >out.log 2>&1 &

‘‘‘

After sketching, a quality control step was implemented to exclude sketches smaller than 20KB, which often represent amplicon or viral data lacking explicit annotation or encountered sketching failures. The remaining sketches were profiled using MetaKSSD, and runs containing fewer than two species were excluded. This process resulted in 382,016 high-quality sketches (Data Availability).

### 4.6 Identifying commonly studied environments

From the metadata of all runs, 66 environments encompassing at least 30 projects were identified (Tab. S4). Thirteen environments were excluded from the final list due to ambiguous annotation or duplication: Homo_sapiens, Mus_musculus, bioreactor_metagenome, feces_metagenome, food_metagenome, freshwater_metagenome, gut_metagenome, human_metagenome, insect_metagenome, marine_metagenome, sediment_metagenome, synthetic_metagenome, and viral_metagenome, resulting in 53 commonly studied metagenomic environments (Tab. S5).

### 4.7 Controlling for suspicious species

To rank species by the number of environments in which they were present, suspicious findings were controlled as follows: a species was considered present in an environment only if it was detected in more than 10% of the projects from that environment, and a species was considered present in a project only if it was detected in more than 10% of the samples from that project.

### 4.8 T-SNE analysis

Projects encompassing at least 30 samples and originating from environments that included at least two different projects with at least 30 samples were first selected. These projects were then manually checked to ensure that the project descriptions were consistent with their “Scientific_name” in the SRA metadata. Projects with ambiguous descriptions of environments or spanning multiple environments were excluded. Finally, 151 projects were selected, encompassing 30,592 samples across 17 different environments (Tab. S7). The MetaKSSD (L3K11S18) profiles were consolidated into a species-by-sample matrix. For samples with multiple runs, only one run was chosen at random. The t-SNE^70^ analysis was performed using the ‘t-SNE’ method from the ‘manifold’ module in the Python ‘sklearn’ package^71^, with parameters perplexity = 30, init = “pca”, metric = “l1”, and n_iter = 80000. The t-SNE results were visualized using the R package ‘ggplot2’^72^.

### 4.9 Construction of abundance vector database, abundance vector searching, and clustering

Each of the 382,016 metagenomic sketches (Methods section 4.5) was profiled, encoded as a sparse abundance vector, and saved as a binary file (*.abv) using the following command:

‘‘‘

metakssd composite -r <markerdb_L3K11> -q <metagenome sketch> -b -o <markerdb_L3K11/abundance_Vec>

‘‘‘

Here, the output path for the abundance vector, “abundance_Vec”, was placed under the MarkerDB directory “markerdb_L3K11” because the construction of the abundance vector database depends on the MarkerDB. Subsequently, all the *.abv files were indexed, and the abundance vector database was created using the command:

‘‘‘

metakssd composite -r <markerdb_L3K11> -i

‘‘‘

To quantify the distance or similarity between two abundance vectors ***a*** = (*a_1_*, *a_2_*, …, *a_n_*)*^T^* and ***b*** = (*b_1_*, *b_2_*, …, *b_n_*)*^T^*, the normalized L1 norm distance and the cosine similarity were employed:

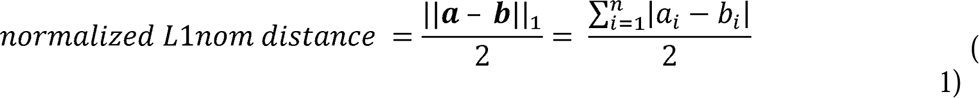

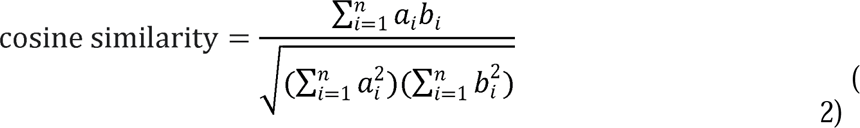

where *n* represented the total number of species in the MarkerDB. To retrieve abundance vectors similar to an input abundance vector (e.g., “input.abv”) from the database, the following command was executed:

‘‘‘

metakssd composite -r <markerdb_L3K11> -s<0 or 1> <input.abv>

‘‘‘

Here, the options -s0 and -s1 specified searching based on normalized L1 norm distance and cosine similarity, respectively.

For abundance vector clustering, the multidimensional scaling (MDS) method^73^ was employed using pairwise normalized L1 norm distance (Eq. 1) or cosine distance as input. Cosine distance can be derived from cosine similarity as follows:

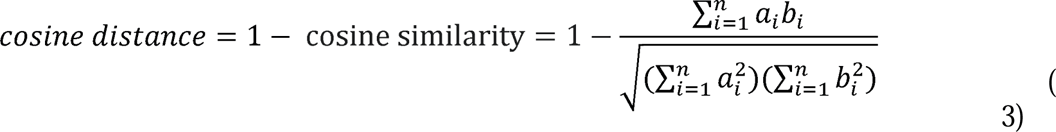

## 5. Data Availability

- **MarkerDB of MetaKSSD (L3K11):** https://zenodo.org/records/11437234/files/markerdb.L3K11_gtdb_r214.tar.gz
- **Abundance vector database of MetaKSSD (L3K11):** https://zenodo.org/records/11437234/files/markerdb.abvdb231227.L3K11_gtdb_r214.tar.gz
- **All metagenome sketches (L3K11, as of 20231227):** Available through zenodo records:10609030,10614425,10614597 and 10676887.
- **The MetaKSSD profiles for all sketched runs:** Available through zenodo record: 11345411
- **Four CAMI2 benchmark datasets:** ‘Mouse_gut’ dataset is available at https://frl.publisso.de/data/frl:6421672/dataset/; ‘Rhizosphere’ ‘Marine’, and ‘Strain_madness’ datasets are available at https://frl.publisso.de/data/frl:6425521/.
- **Previous CAMI2 results:** Available from CAMI2 GitHub repositories: https://cami-challenge.github.io/OPAL/cami_ii_mg/ and https://github.com/CAMI-challenge/second_challenge_evaluation/tree/master/profiling.
- **The OPAL results on the five datasets for all profilers benchmarked in this study:** Available at: yhg926.github.io/KSSD2/OPAL
- **Microbiome-phenotype association study:** The stool microbiome WGS data from the 368 Chinese individuals of BGInature2012 cohort are available under NCBI accession SRA045646. The related metadata is available from the supplementary tables of the source paper^44^.

## 6. Code Availability

- **MetaKSSD (Linux):** https://github.com/yhg926/MetaKSSD
- **MetaKSSD Clients (Mac OS, see <u>tutorial video</u>):** http://www.genomesketchub.com/download/MetaKSSD_Mac.zip
- **MetaKSSD Clients (Windows OS, see <u>tutorial video</u>):** http://www.genomesketchub.com/download/MetaKSSD_Windows.exe

## 7. Supplemental information

Supplemental information includes 10 tables and 11 figures.

## 9. Acknowledgments

This work was supported by 2023 Start-up Funds for Talent at Basic Research Institutions - Yi Huiguang’s Team, (JCKY2023-30); 2023 Basic Research Institution Task 4 - Yi Huiguang (JCKY-ZDKY202304-10); Outbound Postdoctoral Research Funding in Shenzhen (SZS21001); Outbound Postdoctoral Research Funding in Dapeng New District (SDP21029) and Provincial Laboratory Special Start-up Funds - Yi Huiguang’s Team (SSZXQD006).

## 10. Contributions

H.Y. invented the method, developed the software, performed the analyses, wrote the manuscript, and supervised this study. X.L. and Q.C. tested the software and assisted with the analyses.

