## Supplemental Figures for "MetaKSSD: Boosting the Scalability of Reference Taxonomic Marker Database and the Performance of Metagenomic Profiling Using Sketch Operations"

**Supplementary Figures**

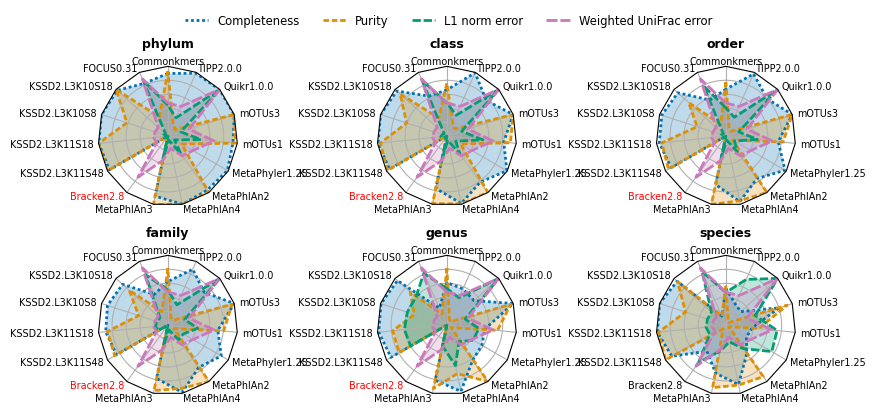

Figure S1 Performance of different profilers across all taxonomic ranks on the mouse gut dataset.

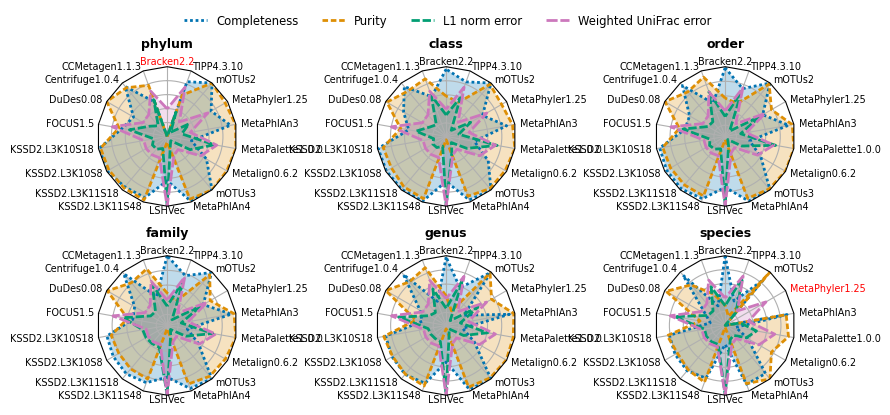

Figure S2 Performance of different profilers across all taxonomic ranks on the marine dataset.

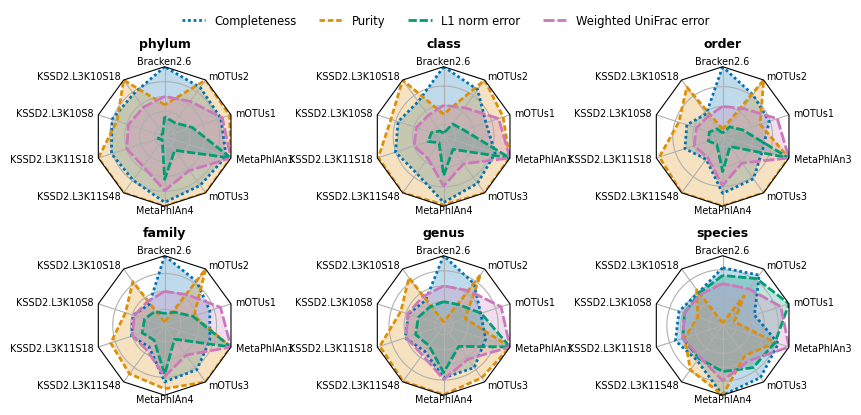

Figure S3 Performance of different profilers across all taxonomic ranks on the rhizosphere dataset.

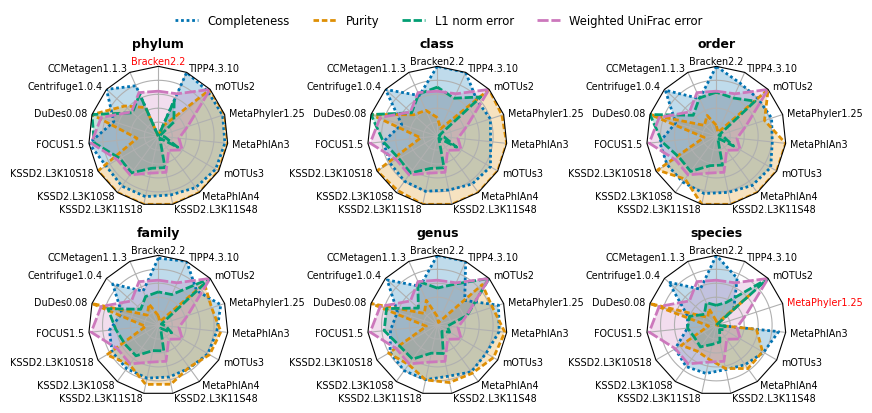

Figure S4 Performance of different profilers across all taxonomic ranks on the strain madness dataset.

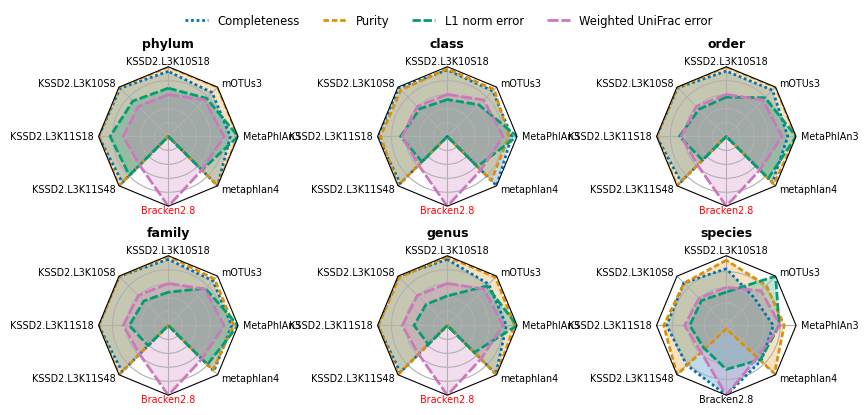

Figure S5 Performance of different profilers across all taxonomic ranks on the [New_released](https://yhg926.github.io/KSSD2/OPAL/new_released)

dataset.

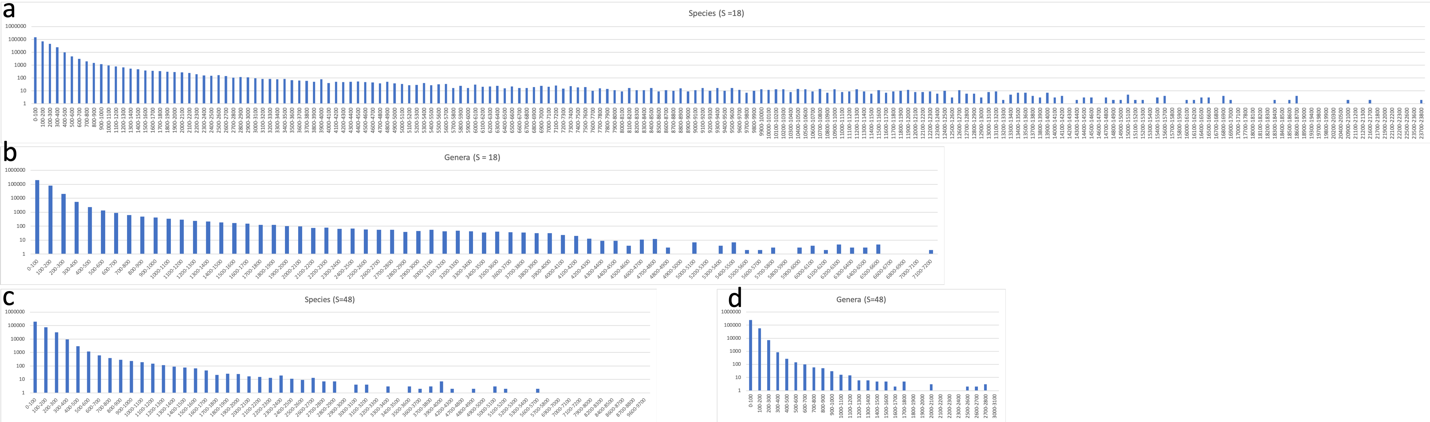

Figure S6. Distribution of Samples Based on the Number of Detected Species and Genera (Grouped in Bins of 100). The summaries were generated based on all the 382,016 metagenome runs profiled by MetaKSSD (L3K11, with MarkerDB constructed from GTDBr214) (see Data availability for all profiles; see Table S*1 for runs information). The taxa (species or genera) number for each run was determined from its raw profile by thresholding the MarkerDB overlapped *k*-mer number (denoted as *S*, the third column) at 18 or 48. And taxa number of a sample is calculated as the average of the taxa numbers of runs from this sample.

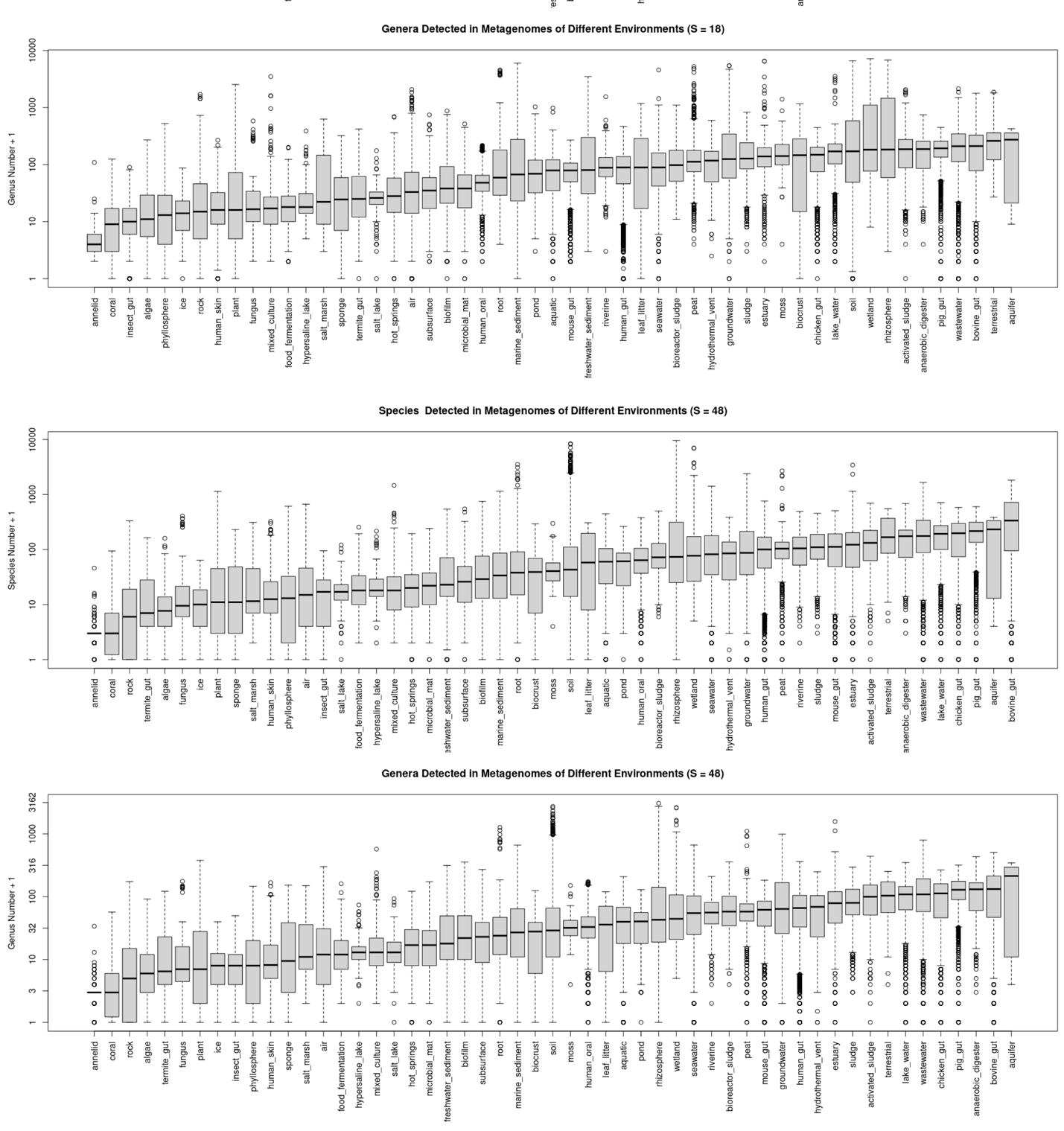

**Fig. S7 Number of Species and Genera Detected in Samples of 53 Commonly Studied Environments.**  The taxa (species or genera) number for each run was determined from its raw profile by thresholding the MarkerDB overlapped *k*-mer number (denoted as *S*, the third column) at 18 or 48. Each data point represents a sample rather than a run, with the value representing the average of taxa numbers of all runs from this sample (referred to Tab. S4 for all runs and samples for these environments).

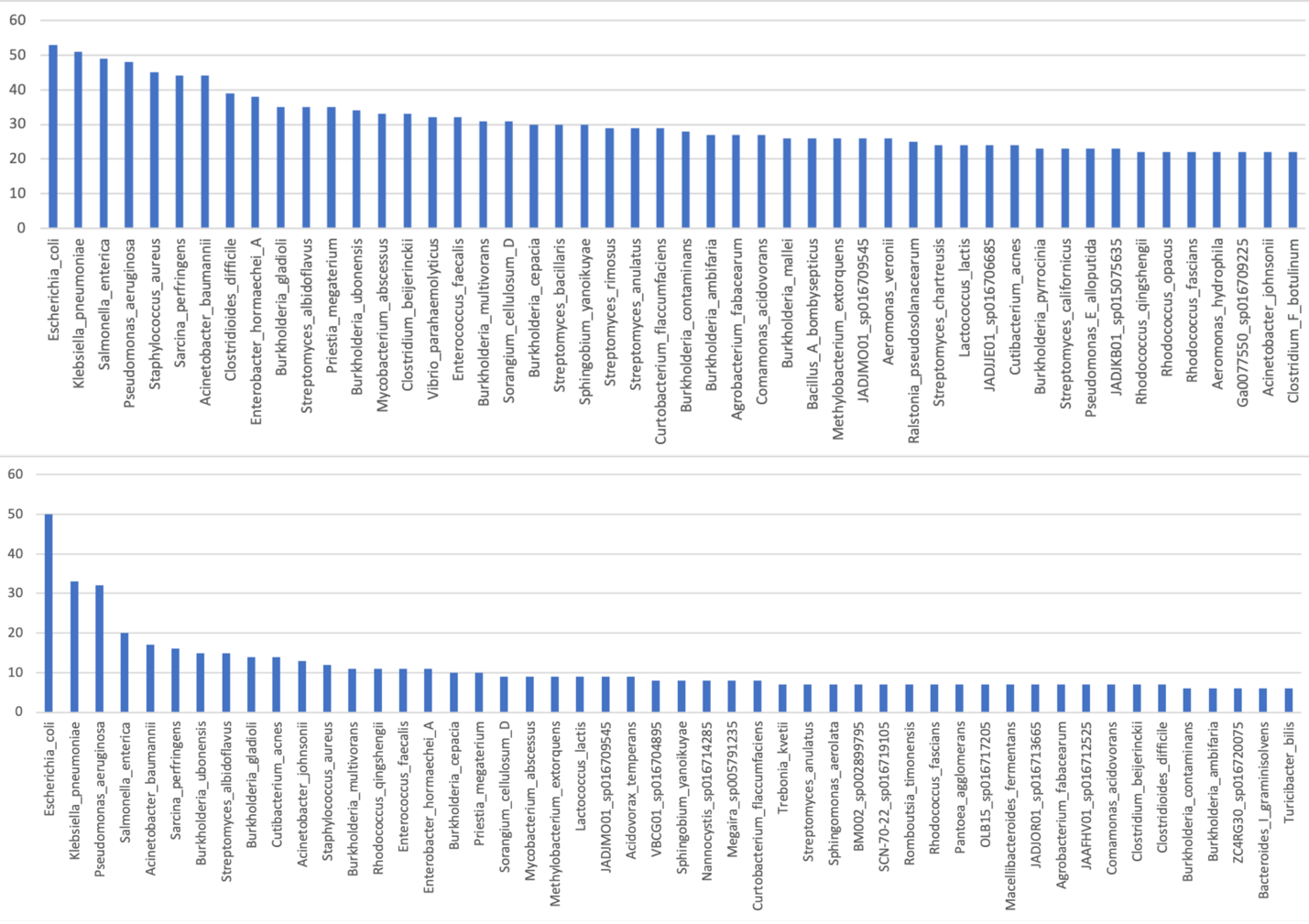

Figure S8 Top 50 Most Commonly Seen Species.

All species detected by MetaKSSD (L3K11), after controlling for suspicious findings (see Methods), were ranked by the number of environments in which they are present. Top panel was summarized from MetaKSSD’s raw profiles thresholding the MarkerDB overlapped *k*-mer number *S*=18, Bottom at *S*=48.

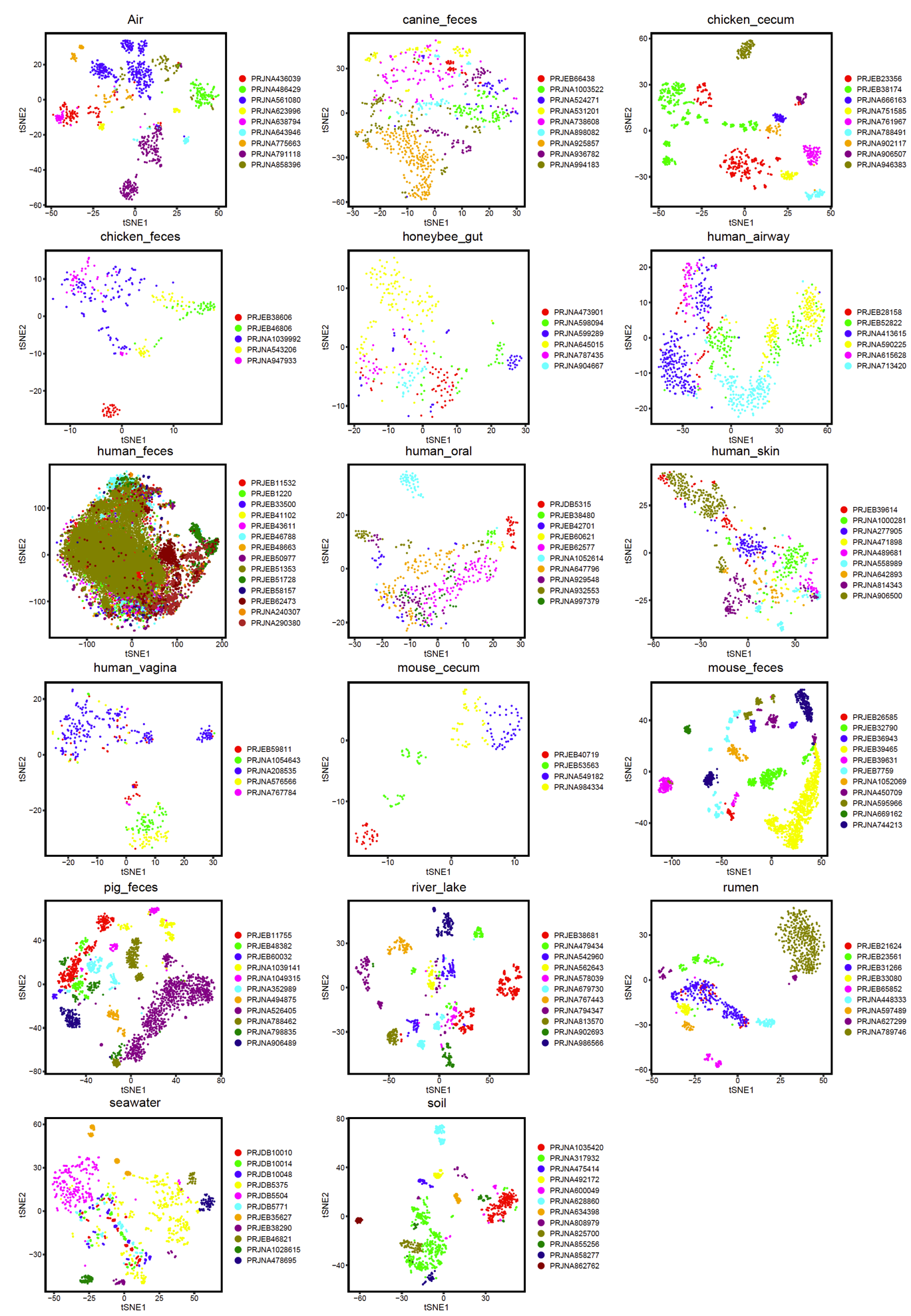

Figure S9 T-SNE analysis of 151 well-characterized projects from 17 environments (Table S7) showed that the sub-clusters within an environment were clearly grouped by projects.

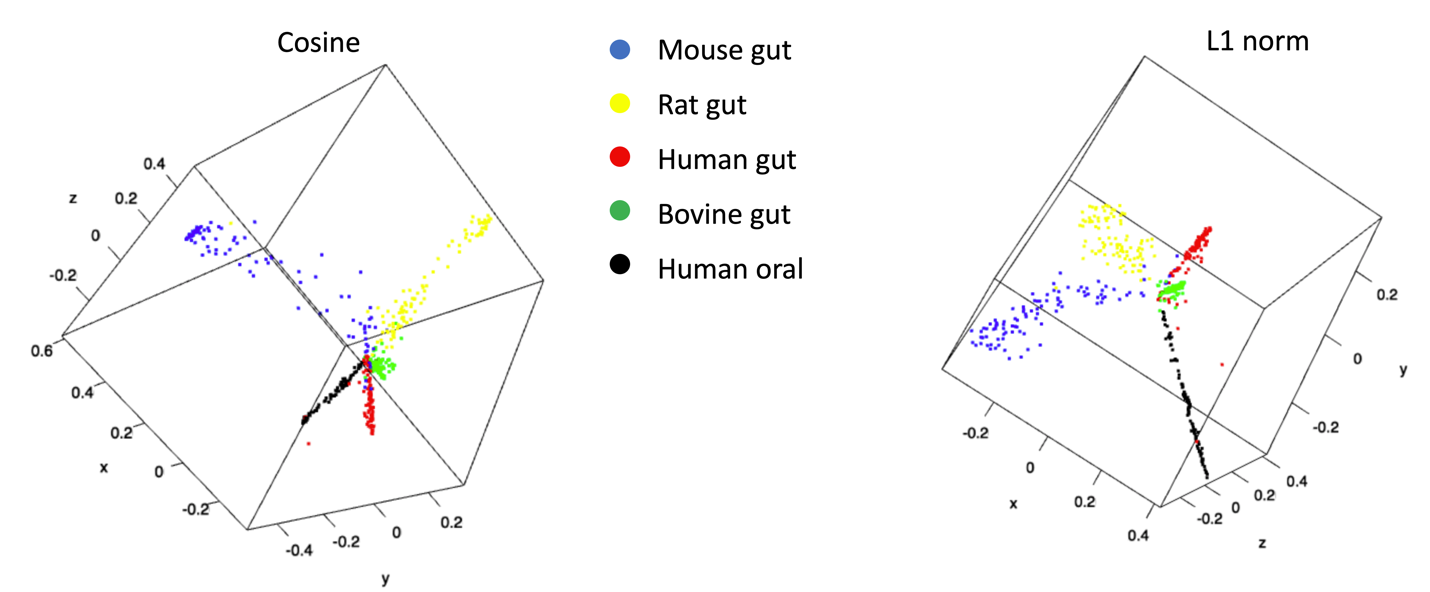

Figure S10. Both L1 norm distance and cosine distances can effectively group samples by their originating environments using the Multiple Dimensional Scaling.

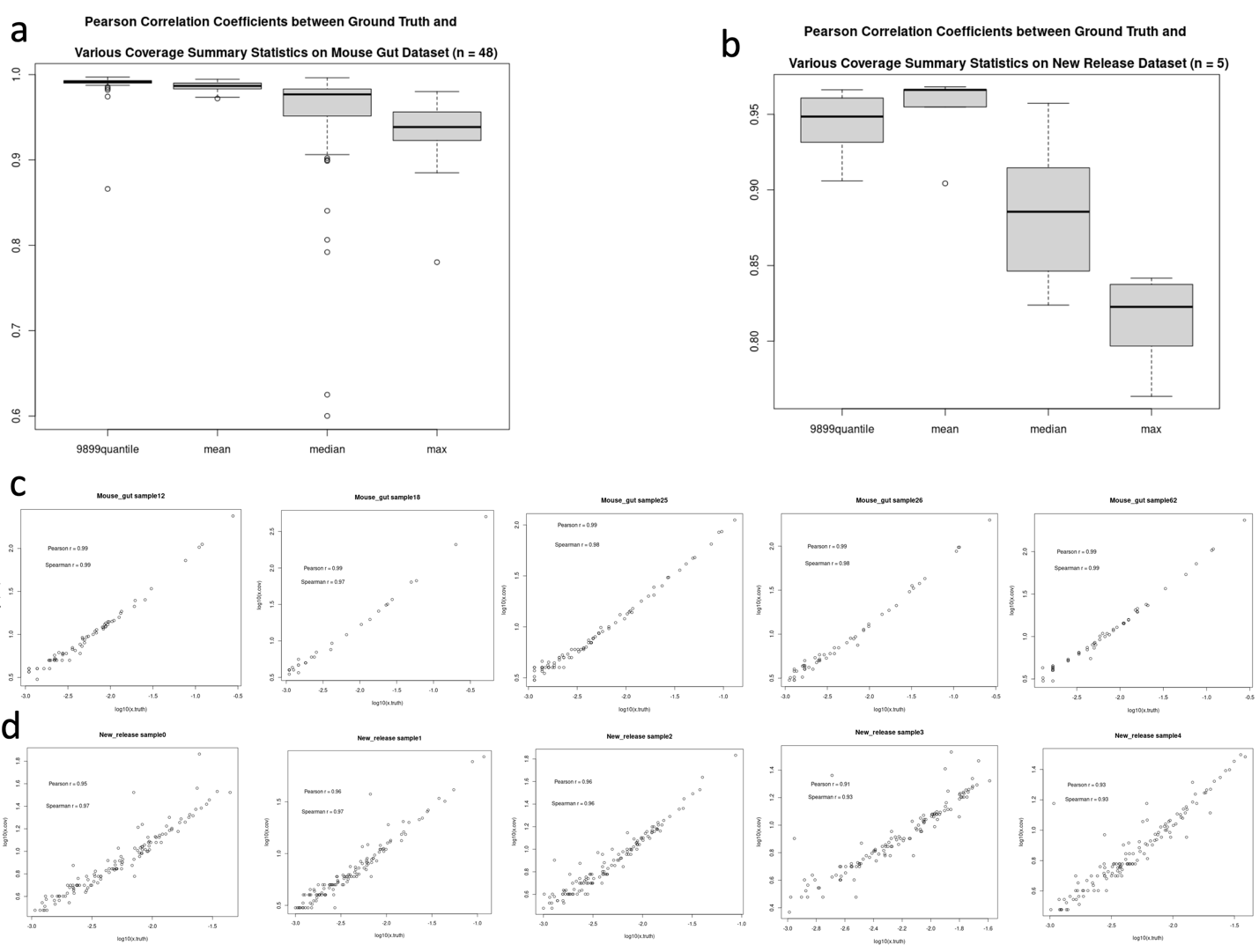

Fig. S11: Performance of Different Summary Statistics of *k*-mer counts. (a) and (b) The 9899quantile represents the mean of the 98^th^ and 99^th^ percentile of *k*-mer counts overlapped between a sample sketch and the MarkerDB; mean denotes the average of overlapped *k*-mer counts; median indicates the median of overlapped *k*-mer counts; max signifies the maximum overlapped *k*-mer count. (c) and (d) Scatter plots comparing truth relative abundance versus 9899quantile *k*-mer counts in randomly selected five samples from the ‘Mouse_gut’ dataset (c) and ‘New_released’ dataset (d). Both the relative abundance and 9899quantile *k*-mer counts were log-transformed, and Pearson and Spearman correlation coefficients (r) were calculated for each plot. To control for false discovery, a threshold was set for the minimal number of overlapped *k*-mers (>18) and minimal species relative abundance (>0.1%).
